# SGLT2 transcriptomic expression atlas supports a kidney-centric role for empagliflozin’s benefits in heart failure

**DOI:** 10.1101/2023.07.03.547550

**Authors:** Omar Mourad, Shabana Vohra, Sara S Nunes

## Abstract

Sodium-glucose cotransporter 2 inhibitors (SGLT2i), such as empagliflozin, have shown remarkable benefits in reducing cardiovascular events and mortality in patients with heart failure (HF) irrespective of diabetic status. Because of the magnitude of the benefits and broad application in both HF with reduced and preserved ejection fraction (EF), there have been concerted efforts to identify a mechanism for the observed benefits. One hypothesis is that SGLT2i act directly on the heart. Given empagliflozin’s high specificity to SGLT2, we reasoned that SGLT2 expression would be a requirement for cells to respond to treatment. Here, we present a comprehensive transcriptomic analysis of *SLC5A2*, which encodes SGLT2, at the single cell level in multiple datasets, confirming *SLC5A2* expression in a subset of kidney epithelial cells but no meaningful expression in other cell types. This was true irrespective of developmental stage, disease state, sequencing method or depth, and species. Our findings support a kidney-centric role for the cardiovascular improvements reported in patients treated with SGLT2i.

---

Recent clinical trials have demonstrated the benefits of SGLT2i, or “gliflozins”, in reducing cardiovascular events and mortality in HF patients irrespective of EF. Notably, these cardioprotective benefits are absent in other glucose lowering drugs and are seen even in nondiabetic patients (1) and thus cannot be explained by the amelioration of hyperglycemia. Moreover, the cardioprotective effects of SGLT2i are extremely rapid, becoming apparent within weeks of treatment (1). Here, we examined the expression of *SLC5A2* at the single cell level, with a focus on cells in the cardiovascular system.

Analysis of the Tabula Sapiens, a human single-cell transcriptomic atlas (2), revealed a cluster of *SLC5A2*-positive cells within the kidney (Figure 1A). Detailed analysis of the kidney revealed that *SLC5A2* is expressed within a subset of epithelial cells (Figure 1B), confirming previous reports on SGLT2 expression in the proximal tubule. In addition, *SLC5A1*, which encodes for SGLT1, an SGLT isoform, was not expressed in kidney cells as expected (Figure 1B).

**Figure 1.**
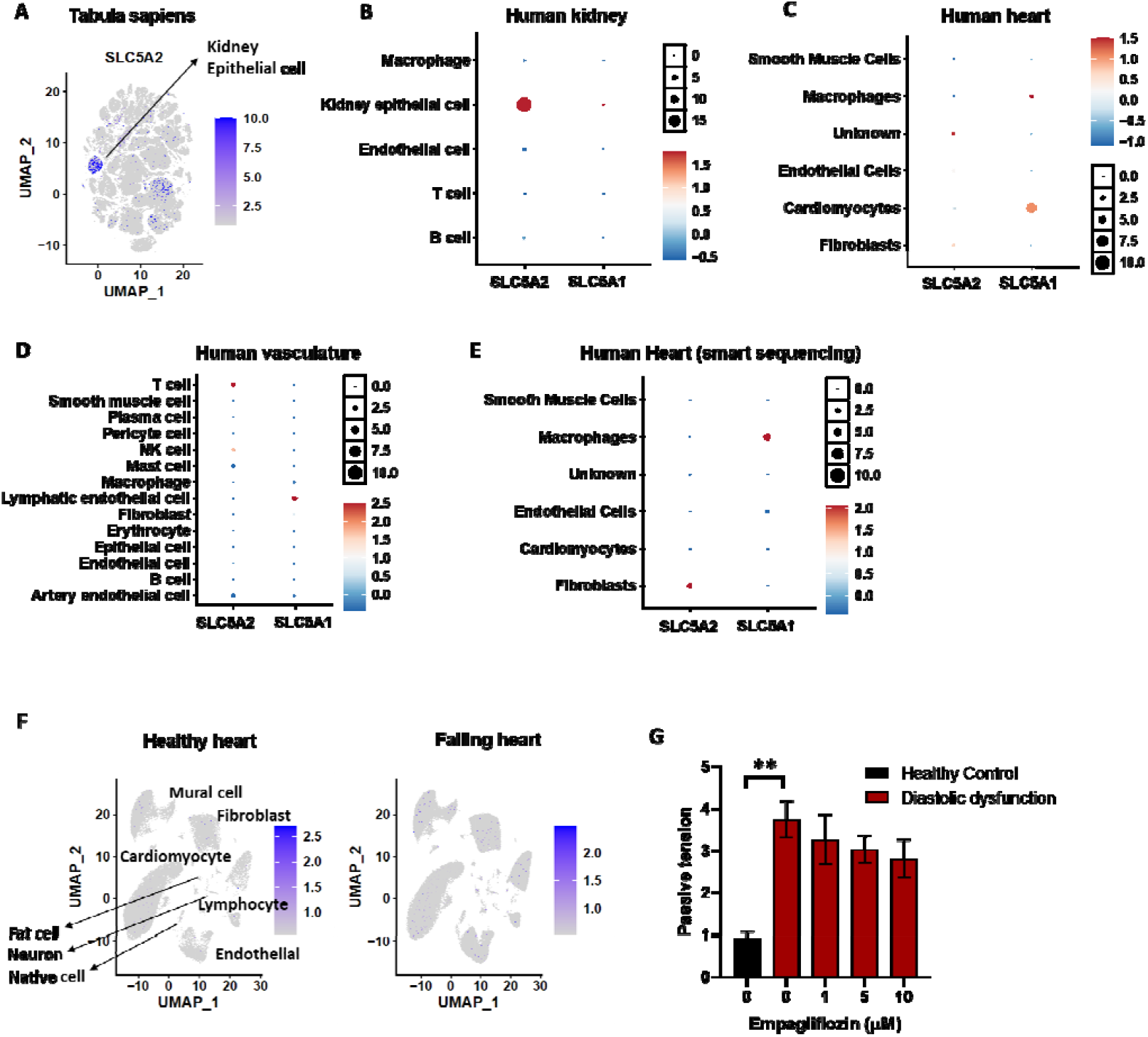
Expression of SLC5A2 in single cell human transcriptomic atlas and disease datasets. **(A)** Feature plot showing the expression of *SLC5A2* in Tabula Sapiens. **(B)** Dot plot depicting the expression and percentage of cells expressing *SLC5A2* and *SLC5A1* in each cell cluster in the human kidney. **(C)** Expression of *SLC5A2* and *SLC5A1* in different cell types in the human heart. **(D)** Dot plot showing the expression of *SLC5A2* and *SLC5A1* in large vessels. **(E)** Dot plot displaying the expression of *SLC5A2* and *SLC5A1* across six human heart clusters from datasets generated by deep-sequencing. **(F)** Feature plot displaying the expression of *SLC5A2* in the single cell transcriptome of normal and failing nonischemic human hearts. **(G)** Treatment of a heart-on-a-chip model of diastolic dysfunction with empagliflozin does not improve function (mean ± SEM; one-way ANOVA Bonferroni’s multiple comparison test. ** P < 0.01).

Given its benefits in HF, we analyzed heart cells which showed a lack of *SLC5A2* expression (Figure 1C). In contrast, *SCL5A1* was expressed in a subset of cardiomyocytes. Analysis of large blood vessels demonstrated that no cells expressed either *SLC5A2* or *SCL5A1* (Figure 1D). One possibility is that the expression levels for *SLC5A2* are low and therefore not detected by traditional single cell transcriptomic methods. Therefore, we analysed deep-sequencing (Smart-seq) data from the Tabula sapiens and confirmed the lack of meaningful *SLC5A2* expression in heart cells (Figure 1E).

Next, we reasoned that SGLT2 expression may be induced in disease and therefore analyzed *SLC5A2* levels in HF datasets. Analysis of transcriptomic data from nonischemic failing and healthy hearts showed that there was no meaningful expression of *SLC5A2* in any of the cell types (Figure 1F) (3). Analysis of a myocardial infarction dataset (4) also revealed the lack of *SLC5A2* expression and comparable *SLC5A1* expression in cardiomyocytes in healthy and infarcted hearts (Figure S1B), demonstrating that cardiac *SLC5A2*/SGLT2 expression is not induced in in these relevant disease states. This indicates that the cardioprotective effects of empagliflozin and other SGLT2i are not due to the direct action in the cells in the cardiovascular system.

Analysis of a human fetal dataset (5) to assess if *SLC5A2* expression may be temporally regulated based on life stage confirmed the lack of expression in cardiac cells in the fetus (Figure S1D). Analysis of the Tabula Muris atlas (6) revealed that *SLC5A2* was only detected in kidney epithelial cells, showing conserved expression patterns across species (Figure S1E).

Moreover, contrary to previous reports in human stem cell-derived cardiomyocytes (hPSC-CMs) that showed improvements in function with empagliflozin treatment (7), we found no functional benefit of empagliflozin (Figure 1G) in a heart-on-a-chip model (8) of diastolic dysfunction that recapitulates hallmarks of diastolic dysfunction (pathological cardiomyocyte hypertrophy, diffuse fibrosis, and increased passive tension compared to healthy tissues) generated by treating cardiac microtissues composed of hPSC-CMs and adult human ventricular fibroblasts with endothelin-1 and transforming growth factor beta-1.

In summary, we have presented a comprehensive transcriptomic analysis at the single cell level showing that there is no prominent expression of *SLC5A2* other than in the kidney epithelial cells, irrespective of developmental stage, disease state, sequencing method or depth, and species. Additionally, no functional benefit was observed with empagliflozin treatment of a diastolic dysfunction heart-on-a-chip model. Given that empagliflozin’s affinity for SGLT1 is >2500-fold lower, we can rule out that the observed benefits in HF are due to interaction with SGLT1. We conclude that the cardioprotective benefits of SGLT2 inhibition are thus likely due to the indirect renal effects of this class of drug. Improved renal function and cardiorenal physiology could ameliorate cardiovascular health by a combination of factors such as natriuresis, reduction in cardiac interstitial edema, improved cardiac bioenergetics, reduced preload and afterload and by reducing left ventricle wall shear stress. Further studies should be aimed at identifying the exact nature of the renal-induced cardiovascular benefits resulting from SGLT2i treatment.

