## Supplemental Figure 1 for "SGLT2 transcriptomic expression atlas supports a kidney-centric role for empagliflozin’s benefits in heart failure"

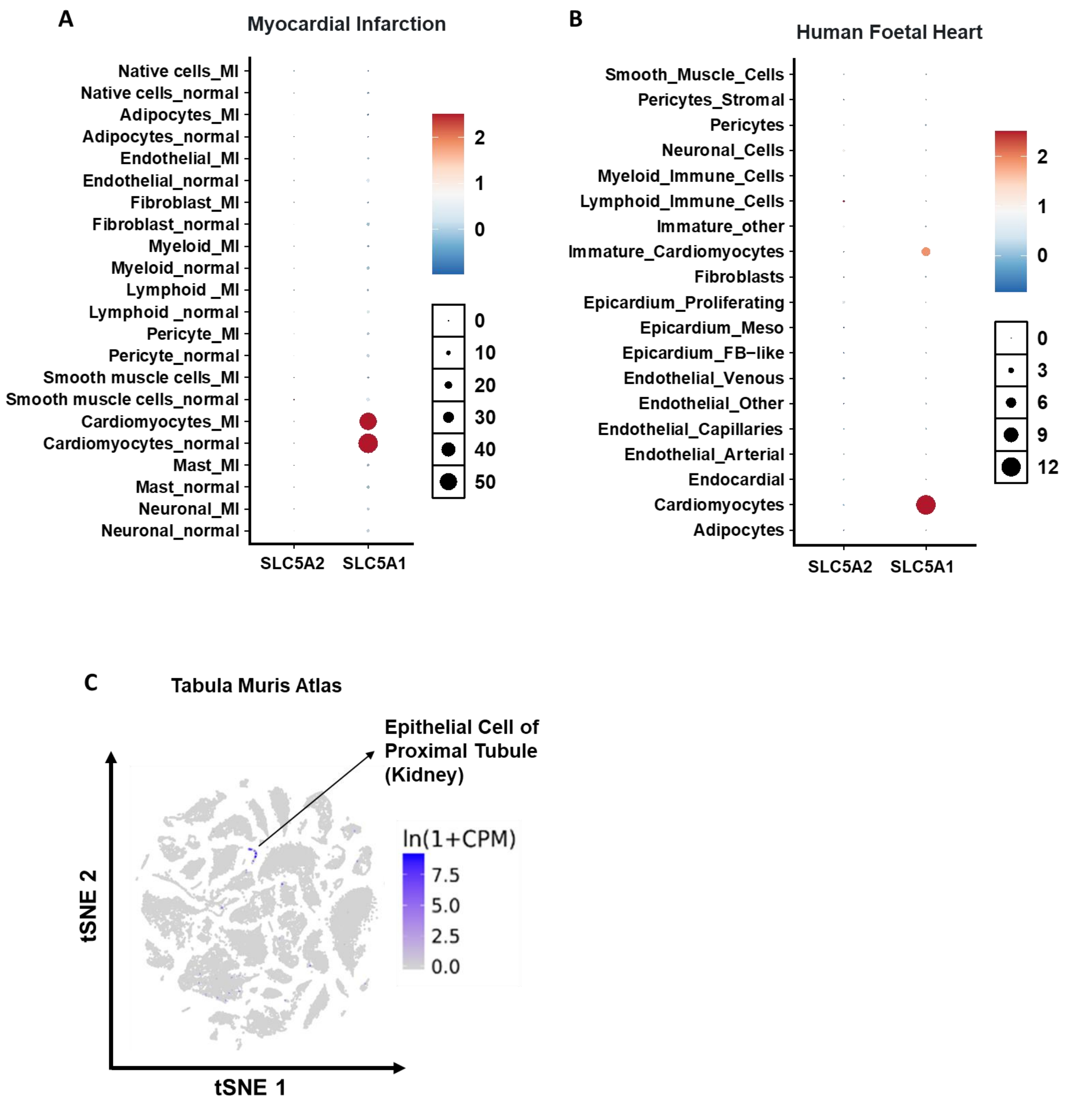


**Supplemental Figure 1. (A)** Expression of *SLC5A2* and *SLC5A1* in myocardium from patients with myocardial infarction and healthy controls. **(B)** Expression of *SLC5A2* and *SLC5A1* across different cell types in human fetal heart. The dataset contains single-cell RNA sequencing of dissociated cells from 7 fetal hearts between the ages of 8 and 12 weeks post-conception. **(C)** Feature plot showing the expression of SLC5A2 gene in Tabula Muris atlas, a compendium of single cell transcriptome data from mice containing nearly 100K cells from 20 organs and tissues.
